# Toward fast and accurate SNP genotyping from whole genome sequencing data for bedside diagnostics

**DOI:** 10.1101/239871

**Authors:** Chen Sun, Paul Medvedev

## Abstract

**Motivation:** Genotyping a set of variants from a database is an important step for identifying known genetic traits and disease related variants within an individual. The growing size of variant databases as well as the high depth of sequencing data pose an efficiency challenge. In clinical applications, where time is crucial, alignment-based methods are often not fast enough. To fill the gap, Shajii et al. (2016) propose LAVA, an alignment-free genotyping method which is able to more quickly genotype SNPs; however, there remains large room for improvements in running time and accuracy.

**Results:** We present the VarGeno method for SNP genotyping from Illumina whole genome sequencing data. VarGeno builds upon LAVA by improving the speed of *k*-mer querying as well as the accuracy of the genotyping strategy. We evaluate VarGeno on several read datasets using different genotyping SNP lists. VarGeno performs 7-13 times faster than LAVA with similar memory usage, while improving accuracy.

**Availability:** VarGeno is freely available at: https://github.com/medvedevgroup/vargeno.

## 1 Introduction

Given a set of target genetic variants, the problem of variant genotyping is to report which variants an individual possesses (Luikart et al. 2003; Shajii et al. 2016; Syvänen 2005). Single nucleotide polymorphism (SNP) genotyping has been widely used in human disease-related research such as genome wide association studies (Hirschhorn and Daly 2005). The approaches to SNP genotyping can be roughly divided into three categories: microarray methods, sequencing alignment-based methods, and alignment-free methods.

The first approach uses SNP arrays (Pastinen et al. 2000). SNP arrays are based on the hybridization of fragmented, single-stranded, target DNA, labelled with fluorescent dyes, to arrays containing immobilized allele-specific oligonucleotide probes (LaFramboise 2009). SNP arrays are fast and inexpensive; however, they can only hold a limited number of probes: the state-of-the-art Affymetrix genome-wide SNP array 6.0 has only 906,000 SNP probes, compared with 31,565,214 known common SNPs in dbSNP (build 150). They also are more narrowly applicable than sequencing when additional analyses are desired.

The second approach is based on high-throughput whole genome sequencing and read alignment. In a standard pipeline using this method, sequencing reads are first aligned to the reference genome. The alignment results are then used as input for genotyping tools such as SAMtools mpileup (Li et al. 2009), or Freebayes (Garrison and Marth 2012), or GATK HaplotypeCaller (Depristo et al. 2011; McKenna et al. 2010). The limitation of this direction is that it requires a lot of time in alignment. This limitation becomes especially crucial in clinical applications, where bedside genotyping of disease-related SNPs may become common in the future.

The third approach is based on high-throughput whole genome sequencing followed by an alignment-free sequence comparison (Vinga and Almeida 2003). Alignment-free methods save compute time and memory by avoiding the cost of full-scale alignment. Recent alignment-free ideas that have made significant application improvements include pseudo-alignment (Bray et al. 2016), lightweight alignment (Patro et al. 2017), and quasi-mapping (Srivastava et al. 2016). Simultaneously, an alignment-free approach has been applied to SNP genotyping by Shajii et al. (2016). They introduce a SNP genotyping tool named LAVA, which builds an index from known SNPs (e.g. dbSNP) and then uses approximate k-mer matching to genotype the donor from sequencing data. LAVA is reported to perform 4-7 times faster than a standard alignment-based genotyping pipeline, while achieving comparable accuracy.

In this paper, we present a data structure for indexing and querying k-mers that is designed for variant genotyping. Our data structure builds on the core data structure of Shajii et al. (2016) but makes use of a Bloom filter and a linear scanning approach. A Bloom filter is a space efficient data structure for improving scalability (Bloom 1970; Broder and Mitzenmacher 2004) that has been widely used in the context of indexing, compressing and searching whole genome datasets (Rozov, Shamir, and Halperin 2014) and large sequence databases (Solomon and Kingsford 2016, 2017; Sun et al. 2017). Furthermore, we incorporate our data structure into a genotyping framework similar to LAVA, but using quality values and a modified mapping criteria to improve speed and accuracy. Finally, we evaluate our approach on several datasets, compare to existing tools, and evaluate the role of our parameters.

## 2 Definitions

A *k*-mer is a sting of length *k* over the four letters DNA alphabet. A *k*-mer can be naturally encoded in *2k* bits. Given a parameter r, we can divide the bits into the upper *r* bits and the lower 2*k* − *r* bits. The Hamming neighborhood of distance 1 for a *k*-mer K, denoted by N(K), is the set of all *k*-mers with a Hamming distance at most 1 to K. We refer to N(K) as the *neighborhood* of K, for short. Notice that K ∈ N(K) and |N(K)| = 3*k* + 1. N(K) can be partitioned into three subsets: 1) the original *k*-mer K, 2) the *upper neighborhood* of K, which is the set of *k*-mers whose encoding differs with K in the upper *r* bits, and 3) the *lower neighborhood* of K, which is the set of *k*-mers whose encoding differs with K in the lower 2*k* − *r* bits.

A Bloom filter for representing a set *S* of n elements is a bitvector of size *m* and *p* independent hash functions *h*_1_, *h*_2_, …, *h_p_*. Each hash function maps an element to a random integer uniformly between 0 and *m* − 1. The bitvector is initialized to an array of zeros. For each element *x* ∈ *S*, the bits *h_i_*(*x*) of the bitvector are set to 1 for 1 ≤ *i* ≤ *p*. To check if an item *y* is in *S*, we check whether all *h_i_*(*y*) are set to 1. If not, then *y* is not a member of *S*. Otherwise, *y* is a member of *S* with a small false positive rate (Broder and Mitzenmacher 2004).

## 3 Methods

Our method uses the same framework as Shajii et al. (2016) and consists of two main ingredients. The first is a data structure to solve the following problem. We are given a set of *k*-mers *S*, with satellite data associated with every *k*-mer. Then, given a *k*-mer *K*, return all the satellite data associated with all the *k*-mers *N*(*K*) that are in *S*. The motivation behind the data structure is that the set *S* contains *k*-mers based on the reference genome, and the query takes a *k*-mer *K* from a donor genome and checks where it matches in the reference. The neighborhood of *K* is used to allow for up to one sequencing error. The second ingredient is a genotyping module that uses the above data structure to call genotypes in a donor. In this section, we describe our data structure (Section 3.1 and 3.2) and genotyping module (Section 3.3).

### 3.1 Indexing data structure description

We choose the value *k* = 32 so that the probability of more than one erroneous nucleotide is low and so that a *k*-mer can be conveniently encoded using a 64-bit integer (Shajii et al. 2016). We also use a parameter *r* to divide an encoded *k*-mer into upper *r* bits and lower 2*k* − *r* bits. In our application, we will use *r* ∈ {24,32}.

#### Index construction

We construct a dictionary *D* from the set *S. D* is an array of <encoded *k*-mer, satellite-data-pointer> tuples, sorted in increasing order of encoded *k*-mers. We also construct a secondary indexing hash table *J* which maps an *r*-bit unsigned integer *u* to the first location in *D* at which there is an encoded *k*-mer whose upper bits are *u*. Finally, we build a Bloom filter *B* where each element is the lower 2*k* − *r* bits of a *k*-mer in *S*.

#### Query algorithm

To query a given *k*-mer *K* and its neighborhood *N*(*K*), we proceed in two steps (illustrated in Figure 1). In the first step, we perform an *upper neighborhood* query, which searches for all the *k*-mers that are in the upper neighborhood of *K*. We first check if the lower 2*k* − *r* bits of *K* exist in *B*. If no, then we abandon the upper neighborhood search. If yes, then for every *k*-mer *K*′ in the upper-neighborhood, we query *J* to find the start and end of the block in *D* with the same upper bits as *K*′. We then do a binary search through this block to find an entry that matches *K*′ if it exists. Note that because *B* contains false positives, there may be no match in *D*.

**Figure 1.**
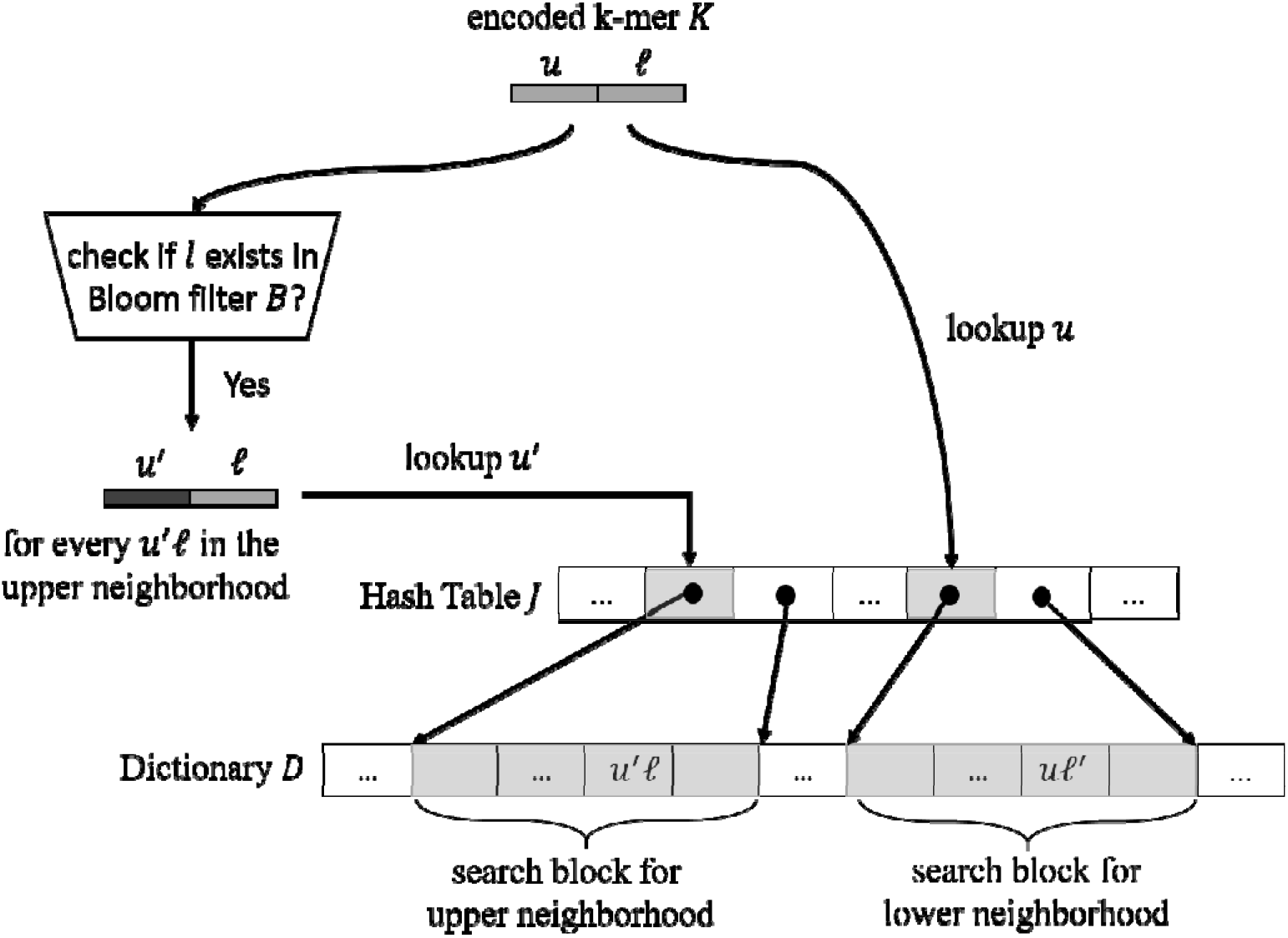
VarGeno’s query algorithm. The encoded *k*-mer *K* can be divided into the upper *r* bits and the lower 2*k* − *r* bits, represented as *u* and *l*. VarGeno starts by checking if *l* exists in Bloom filter *B*, and if yes, performing the upper neighborhood query as follows. It first looks up the upper bits of each *k*-mer in the upper neighborhood of *K* (represented as *u*’) in index *J*. This locates the search block in dictionary *D* for all *k*-mers whose upper bits are it’. Next, VarGeno performs the lower neighborhood query by first looking up *u* in *J*. This locates the block in *D* of k-mers whose upper bits are *u*. In this figure, we will assume that the block is larger than the threshold. VarGeno than scans the block and for each element *uℓ*′ checks its Hamming distance to *uℓ*.

In the second step, we perform a *lower neighborhood* query, which searches for all the *k*-mers that are in the lower neighborhood of *K*. First, we query *j* to find the start and end indices of the block in *D* with the same upper *r* bits as *K*. Then, if the size of the block is larger than a size threshold (given as a parameter *t* to the algorithm), for every *k*-mer in *N*(*K*), we do a binary search to find if it exists in this block. If the size of the block is smaller than *t*, we instead do a linear scan of the block and, for every element, compute its Hamming distance to *K*. A hit is reported if the distance is at most 1. The Hamming distance computation is done using a fast bitwise routine which also identifies where the differing position is (see Supplementary Information).

### 3.2 Indexing data structure performance

Our data structure is based on the data structure of Shajii et al. (2016) with two key differences: the use of the Bloom filter in the upper neighborhood query and the use of a linear scan in the lower neighborhood query. The improvements of our data structure are heuristic in nature and hence a formal analysis did not yield any insights. Here, we argue why these heuristics can improve running time.

For the upper neighborhood query, each *k*-mer in the upper neighborhood of *K* will have different upper bits and hence will require a separate access to *J* and one random access to *D*. The random access to *D* will likely result in a cache miss for every *k*-mer in the upper neighborhood. By using the Bloom filter *B*, we make sure to pay this cost only for *k*-mers that are likely to result in hits.

For the lower neighborhood query, using a linear scan when the block size *t* is small can have a significant improvement on performance. In the worst case, binary search requires 16 * 3 + 1 searches of *O*(log *t*) time each, i.e. 49 * *O*(log *t*) comparisons. A linear scan, on the other hand, requires only *t* calls to the highly optimized Hamming distance routine. In the Results, we investigate which block size threshold yields improved performance.

While binary search is asymptotically faster, its overhead relative to a linear scan makes it slower when the size of the block being searched is small. We found that using *t* = 25 works well in practice.

The following Observation and analysis shows that the number of blocks larger than *t* is small.

#### Observation 1.

Let *n* be the number of distinct *k*-mers stored in the dictionary *D* and let *b* be the number of blocks in *D*. Under the assumption that the encoded *k*-mers are independent from each other, the size of a block in *D* is at least *t* with probability of at most 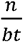.

**Proof.**

Let *x_i_* be the size of block *i* in *D*. Under the independence assumption, *x_i_* follows a Binomial distribution with *n* trials and success probability of 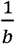. The expected size of block *i* is therefore 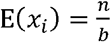. Applying Markov’s inequality (Buot 2006), the probability that *x_i_* is at least *t* is 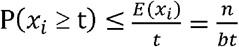.

The assumption that encoded *k*-mers are independent from each other is not true in our application since many of the k-mers overlap. However, we argue that block locations of two encoded k-mers are much less dependent. Even for overlapping *k*-mers, a one nucleotide difference in the higher-order bits will significantly change the encoded k-mer value and, hence, the block location.

For the human genome, the number of *k*-mers n is about 3 billion and *b* is about 2^32^. The probability that a block is larger than *t* = 25 (VarGeno’s default threshold) is less than 0.028. Thus, the Observation estimates that we resort to the binary search method for less than 3% of the blocks.

### 3.3 Genotyping module

We now describe how our genotyping module works and integrates the indexing data structure.

In VarGeno, one index is constructed for all the *k*-mers in the reference genome (using *r* = 32). Another index is constructed with *k*-mers from positions that overlap some SNP from the population SNP database (called the *SNP list*), with the reference allele replaced by alternate allele (using *r* = 24). Then, given reads from a donor genome, VarGeno splits each read into non-overlapping *k*-mers and queries the two indices with them (reverse complements are handled in the implementation but, for the sake of brevity, are not accounted for in our description). However, VarGeno does not explore the whole Hamming neighborhood: if the quality score for a certain position within the query *k*-mer is higher than a threshold, then the neighborhood *k*-mers which differ at this position are skipped and not looked up in the dictionaries. The intuition behind this is that a sequencing error in a position with a high quality score is unlikely.

After querying the *k*-mers from a read, VarGeno determines a single mapping location for the read. A read is considered mapped to a reference genome location if 1) that location has the most hits from the queries, 2) at least two of the hitting *k*-mers originate from different positions in the read, and 3) at least one *k*-mer should be non-modified (i.e. present in the read without a substitution). A read is discarded if more than one location satisfies these criteria.

Once the read’s best matching location on the reference genome is decided, the read is used to support either the reference or alternate allele of SNPs inside the matching location. The information is stored in a pileup table. After processing all reads, the pileup table is then used to compute the most likely genotype for each SNP in the SNP list following the formulas in Shajii et al. (2016). Finally, we note that our genotyping module is similar to Shajii et al. (2016) with the difference that LAVA does not use quality values and only uses the first criterion for determining a mapping location for a read.

## 4 Results

We implemented VarGeno in a C++ package, building on the LAVA code base (Shajii et al. 2016) and code from Sun et al. (2017). VarGeno is freely available at: https://github.com/medvedevgroup/vargeno. VarGeno fixes the number of hash function in the Bloom filter to be one, to reduce the hashing time. All experiments were run on an Intel Xeon CPU with 512 GB of RAM using a single core (at 2.10 GHz). Default parameters were used for all tools, unless otherwise noted.

### 4.1 Experimental setup

Our first dataset is a set of NA12878 reads from Phase 1 of the 1000 Genome Project (1000 Genomes Project Consortium et al. 2012). The reads are 101nt long and the depth of coverage is around 6X. To further benchmark on higher coverage datasets, we used a set of NA12878 reads from Genome in a Bottle Consortium (GIAB) (Zook et al. 2014). The dataset contains reads with length 148, and we randomly selected three subsets of reads with depth of coverage around 15X, 25X and 51X, respectively.

We use two different SNP lists. The *dbSNP-list* contains all common SNPs from dbSNP (11,129,706 SNPs; build 142). The *affy-list* contains 943,192 SNPs that are used by the Affymetrix SNP chip (McCarroll et al. 2008). This is a smaller SNP list with few dense regions (32bp windows with more than one SNP) than the dbSNP-list, and easier to genotype. We used GRCh37/hg19 as the reference sequence.

For validation, we used an up-to-date high-quality genotype annotation generated by GIAB (Zook et al. 2014). The GIAB gold standard contains validated genotype information for NA12878, from 14 sequencing datasets with five sequencing technologies, seven read mappers and three different variant callers. To measure accuracy, we use loci in the SNP list which are also genotyped in the GIAB gold standard (so called high confident regions). Genotypes not reported explicitly are considered as homozygous reference by default, for all genotyping methods.

### 4.2 Comparison against alignment-based discovery pipelines

We compared the performance of VarGeno against two alignment-based discovery pipelines on the 6X dataset and the dbSNP-list (Table 1). The first pipeline, denoted by BWA+mpileup, runs BWA-mem (Li and Durbin 2010), followed by samtools mpileup (Li et al. 2009), followed by ‘bcftools call –gf’ (Narasimhan et al. 2016). The second pipeline, denoted by BWA+GATK-HC ran BWA-mem, followed by the up-to-date best practice suggested by GATK (Depristo et al. 2011; McKenna et al. 2010).

**Table 1.** Performance of VarGeno and LAVA compared to alignment-based approaches on the 6X dataset and the dbSNP list.

| Category | Algorithm | Run time (mins) | Memory usage (GB) | Accuracy (%) |
| --- | --- | --- | --- | --- |
| Alignment-based | BWA+mpileup | 1,800 | 3.2 | 93.46 |
|  | BWA+GATK-HC | 2,400 | 17.0 | 91.80 |
| Alignment-free | LAVA | 315 | 59.7 | 91.90 |
|  | VarGeno | 29 | 60.9 | 93.45 |

Between the alignment-based pipelines, BWA+mpileup performs better than BWA+GATK HC in all aspects, which agrees with the previous reported observation (Shajii et al. 2016). VarGeno is more than 62 times faster with the same level of accuracy as BWA+mpileup, though with substantially more memory use. We explore a lower memory version of VarGeno in Section 4.5.

### 4.3 Comparison against LAVA

Next we compare VarGeno against LAVA using the 6X and higher coverage GIAB datasets on the dbSNP-list (Tables 1 and 2). VarGeno is 7-13 times faster than LAVA on all benchmarked datasets. The memory usage of LAVA and VarGeno is dominated by the size of the indices and is 2% higher for VarGeno then LAVA (Table 1).

**Table 2.** Performance of VarGeno and LAVA on the dbSNP list, with different read depth of coverage. LAVA’s runtime on 51X data is not shown since we could not maintain isolated server conditions long enough (~2 days) to generate an accurate benchmark. The memory usage of LAVA and VarGeno with higher coverage reads is the same as with the 6X reads in Table 1 and is not shown.

| Read Coverage | Algorithm | Run time (mins) | Accuracy (%) |
| --- | --- | --- | --- |
| 15x | LAVA | 820 | 94.54 |
|  | VarGeno | 66 | 96.80 |
| 25x | LAVA | 1,392 | 94.69 |
|  | VarGeno | 108 | 97.20 |
| 51x | LAVA | — | 94.58 |
|  | VarGeno | 196 | 97.34 |

VarGeno’s accuracy is 2-3 percentage points higher than LAVA’s, due to its use of quality values and modified mapping criteria (Tables 1 and 2). The genotyping accuracy of VarGeno and LAVA increases with coverage but starts to plateau after 15x.

We also tested VarGeno using a different SNP list. Table 3 shows the results for the 6X dataset on the affy-list. The speed-up relative to LAVA is consistent with Tables 1 and 2. However, on this dataset we observe a slight decrease in accuracy with VarGeno’s default parameters. We believe this is due to the fact that affy-list contains less dense regions. However, we note that if VarGeno’s quality threshold parameter is set so that it explores the whole Hamming neighborhood, regardless of quality scores, then its accuracy matches LAVA while still being 46% faster (Table 3, third row).

**Table 3.** Performance of the 6X dataset using the Affymetrix SNP list. The second row shows VarGeno run with the default value for the quality value cutoff (*c* = 23).

|  | Running time (mins) | Memory usage (GB) | Accuracy (%) |
| --- | --- | --- | --- |
| LAVA | 160 | 40.8 | 92.99 |
| VarGeno | 25 | 43.8 | 92.50 |
| VarGeno (c=42) | 87 | 43.8 | 93.07 |

### 4.4 Achieving higher accuracy

With sufficient coverage, VarGeno achieves >97% accuracy on the dbSNP-list (Table 2); however, in medical diagnostic applications an even higher accuracy may be desired. Since increasing the depth of coverage beyond 15x has only a minor effect (Table 2), we looked at alternate ways to improve accuracy. We observed that most of the errors occurred in dense regions. Because of linkage disequilibrium, SNPs are often used as markers for nearby variation. In such cases, the SNP list is unlikely to have many dense regions. To emulate this scenario, we scanned through the dbSNP-list and, for each 32nt genome window that has more than one SNP in it, filtered out all but the first SNP. This resulted in a list of 4,162,639 SNPs (37.4% of dbSNP-list, but more than four times the affy-list). VarGeno’s accuracy on this filtered SNP list is 1.4-1.7 percentage points higher than on the dbSNP-list, reaching 98.75% on the 51X dataset (Table 4).

**Table 4.** Accuracy of VarGeno on the filtered dbSNP-list.

| Read Coverage | Accuracy (%) |
| --- | --- |
| 6x | 95.03 |
| 15x | 98.28 |
| 25x | 98.63 |
| 51x | 98.75 |

### 4.5 Memory-lite version

VarGeno uses around 60 GB of RAM for the dbSNP-list experiments and 44 GB for the affy-list, most of which is used to store the *k*-mer indices. To decrease memory usage, we use an idea from Shajii et al. (2016) to provide a memory-lite version called VarGeno Lite. Instead of including every *k*-mer from the reference genome in the reference index, we only include *k*-mers that are within one read length range of some SNP in the SNP list. Table 5 shows the results on the 6X dataset, compared with the memory-lite version of LAVA (Shajii et al. 2016).

**Table 5.** Performance of memory-lite algorithms on the 6X dataset.

| SNP list | Algorithm | Running time (mins) | Memory usage (GB) | Accuracy (%) |
| --- | --- | --- | --- | --- |
| dbSNP-list | LAVA Lite | 464 | 33.0 | 91.62 |
|  | VarGeno Lite | 50 | 35.2 | 93.44 |
| affy-list | LAVA Lite | 210 | 15.2 | 92.96 |
|  | VarGeno Lite | 20 | 15.5 | 92.49 |

The memory-lite version reduces memory by 44% for the dbSNP-list (Table 1) and by 64% for the affy-list (Table 5). For the affy-list, this means that the algorithm can almost be run on a commodity desktop computer with 16 GB RAM. Surprisingly, the accuracy is nearly identical, with differences less than 0.3 percentage points compared to Tables 1 and 3. The running time change is not consistent with respect to the full memory versions, with an increase for the dbSNP-list (Table 1) and a decrease on the affy-list (Table 3). The relative speed advantage of VarGeno to LAVA remains roughly the same in their corresponding memory-lite versions.

### 4.6 Effect of *k*-mer index optimizations

VarGeno’s algorithm can be viewed as the base LAVA algorithm + use of Bloom filter + linear scanning + quality value cutoff + modified mapping criteria. In this section we investigate the contribution of the Bloom filter and linear scan optimizations in isolation, using the 6X dataset with the dbSNP-list. Table 6 shows the result of applying only the Bloom-filter optimization to LAVA’s algorithm. It reduces the run time by 46%, at the expense of only a 2% increase in memory usage.

**Table 6.** The effect of using Bloom filters to accelerate genotyping.

|  | Running time (mins) | Memory usage (GB) |
| --- | --- | --- |
| LAVA | 315 | 59.7 |
| LAVA + Bloom filters ($m = 8n$) | 171 | 60.9 |
| LAVA + Bloom filters ($m = 16n$) | 170 | 62.1 |

We also measure the effect of varying the size of the Bloom filter (denoted by *m*). A larger size decreases the false positive rate and hence the number of unnecessary queries to the dictionaries; a smaller size decreases the memory usage. VarGeno’s default setting is, where is the number of distinct values that are stored in a Bloom filter. This corresponds to a theoretical false positive rate of 0.118 (Broder and Mitzenmacher 2004). We also tried, which corresponds to a theoretical false positive rate of 0.06. Our results indicate that there is not a significant change in running time or memory usage, relative to the totals (Table 6).

We also measured the cache usage improvements by the Bloom filter optimization with the Linux profiler ‘perf.’ LAVA had total cache misses, while there were only (66%) cache misses after the Bloom filter optimization. This result is consistent with our hypothesis that using the Bloom filter reduces run time mainly by reducing the number of cache misses.

Next, we measured the effects of the linear scan optimization. Adding only the linear scan optimization to LAVA resulted in an improvement of 38.5% to the run time. We also measure the running time as a function of different block size thresholds (Figure 2). Performance drastically improves as long as threshold is at least ten. The performance is not substantially impacted by further increasing the threshold, though it does slightly decline again when it the threshold is very large.

**Figure 2.**
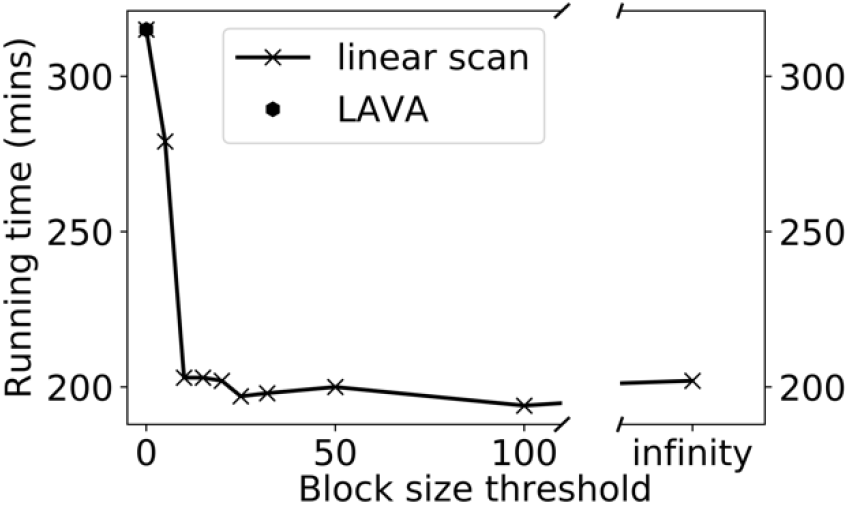
Speed-up due to the linear scan optimization in isolation, as a function of the block size threshold (.A block size threshold of zero is equivalent to what LAVA does. If threshold is infinity, a linear scan is applied to every block.

### 4.7 Effect of the quality value cutoff

We studied the effect that the quality value cutoff optimization has on performance. Recall that VarGeno does not generate neighbors at positions with Phred quality score (Cock et al. 2009) more than some threshold *c*. Figure 3 shows how this parameter affects performance on the 6X dataset with the dbSNP-list.

**Figure 3:**
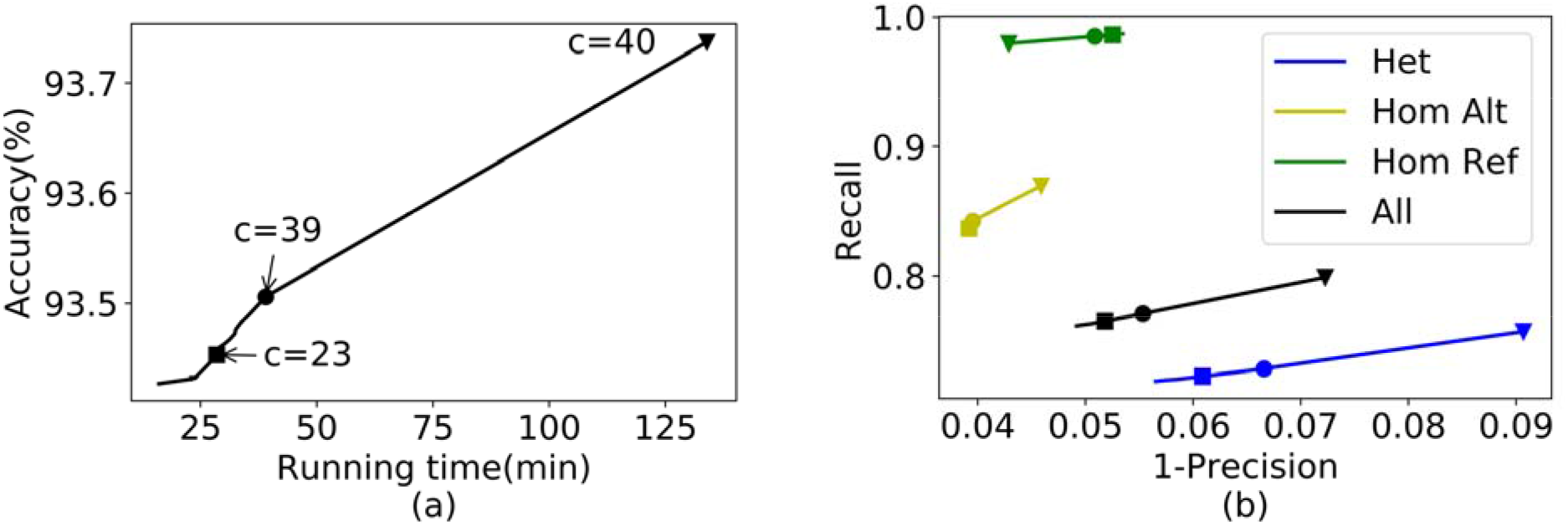
The performance of VarGeno on 6X reads set with the dbSNP-list, under different quality value cutoffs, which are Phred scores in the integer range of [0,42], (a) The accuracy and running time at various cutoffs, (b) The receiver operating characteristic (ROC) curve over different subset of variants in dbSNP-list, with respect to quality value cutoffs. The green line (respectively, the yellow and blue line) represents the subset of variants in dbSNP-list with a homozygous reference (respectively, a homozygous alternate and a heterozygous) genotype in the GIAB gold standard. The black line represents all the variants together. Three representative quality value cutoffs are highlighted using a square (c=23), a circle (c=39) and a triangle (c=40) on each line.

First, we observe a trade-off between running time and accuracy (Figure 3a). The highest accuracy is achieved at (i.e. the highest quality score), which is equivalent to disabling the quality value cutoff and generating all Hamming neighbors. The fastest running time is achieved at, which is equivalent to not exploring any of the Hamming neighborhood. Second, we observe a trade-off between recall and precision (Figure 3b, black curve), with achieving the highest recall and achieving the highest precision. We note that in all cases, VarGeno is faster and more accurate than LAVA. By default, VarGeno uses to achieve a balanced performance.

We further looked at the effect of the quality score separately for loci that are homozygous for the reference allele (according to the GIAB gold standard), heterozygous, or homozygous for the alternate allele (Figure 3b, green, blue, and yellow lines, respectively). Interestingly, the trade-off between recall and precision happens in the opposite direction for loci that are homozygous for the reference allele than it does for all other loci. In other words, using a higher quality value cutoff helps improve the recall of variants (i.e. loci with either an alternate allele present) but decreases the recall of non-variants (i.e. loci with only the reference allele). We observe this affect also when looking at the raw counts of correctly called loci (Table 7).

**Table 7.** Number of correctly reported genotypes by VarGeno based on the genotypes in GIAB gold standard. The first row shows the number of SNPs in dbSNP-list with homozygous reference, heterozygous, and homozygous alternate genotypes in GIAB gold standard. The next three rows show, for three representative quality value cutoffs, the number of correct calls for each subset of the SNP database. For c = 0, the absolute numbers of correct calls are presented. For c = 23 and 42, the relative changes compared to c = 0 are presented.

| Counts in GIAB gold standard |  | Hom Ref: 7,463,731 | Heter: 1,363,475 | Hom Alt: 812,638 |
| --- | --- | --- | --- | --- |
| Correct calls by VarGeno | c = 0 | 7,312,827 | 1,018,751 | 702,699 |
|  | c = 23 | - 6,768 | + 5,337 | + 2,890 |
|  | c = 42 | - 55,300 | + 54,565 | + 30,742 |

## 5 Conclusions

In this paper, we presented VarGeno, an alignment-free SNP genotyping method. We demonstrated that it is more accurate and 7-13 times faster than LAVA, the state-of-the-art alignment-free method. We also compared VarGeno to alignment-based discovery approaches, and, on genotyping, it performs 62 times faster with the same accuracy. VarGeno’s performance advantages are consistent for different SNP lists, such as all the common SNPs in dbSNP (~11 million SNPs) or the SNPs used in an Affymetrix SNP chip (~1 million SNPs). We also demonstrate that even higher accuracy (98.75%) can be achieved by filtering out SNPs from the SNP list that are less than 32nt away from each other.

VarGeno is a streaming algorithm: it can process reads on-the-fly as they come off a sequencer. This is especially useful for variant genotyping scenarios where time is crucial, such as in clinical applications. For instance, in our experiment, VarGeno can genotype 11 million known SNPs from 25x human whole genome sequencing data within 1.8 hours, with accuracy 97.2%. VarGeno can be applied more widely to portable medical devices, if either the genotyping is chromosome specific, or the memory usage can be further reduced for whole genome sequencing data. One possible way to achieve this, at the cost of running time, is to process the reference in separate chunks. Techniques to further reduce memory usage are a future research direction.

## Acknowledgements

This work has been supported in part by NSF awards DBI-1356529, CCF1439057, IIS-1453527 to PM. We would like to thank the anonymous reviewers for their suggestions that led to improving the manuscript.

## Conflict of Interest

none declared.

